# Rapid and reproducible haplotyping of complete mitochondrial genomes using split *k-*mers

**DOI:** 10.1101/2025.03.23.644767

**Authors:** Douglas S. Stuehler, Liliana M. Cano, Michelle Heck

## Abstract

Accurate phylogenetic analysis of mitochondrial haplotypes underpins a wide spectrum of biological inquiry. While multilocus sequence-based phylogenies are standard, inconsistent gene selection and trimming limit network reproducibility and disregard non-coding sequence. To examine alternative approaches, we tested split *k*-mer analysis (SKA) to reassess published mitogenome datasets of one mammalian and two insect species. SKA accurately haplotyped each dataset, improved polymorphism detection in the hemipteran *Diaphorina citri* and aided in the identification of *D. citri* haplotypes associated with the titer of “*Candidatus* Liberibacter asiaticus”, the bacterium associated with citrus greening disease. We present a new mitogenome haplotyping method and script, ska-mtdna.py.

## Background

Accurate phylogenetic analyses inform countless aspects of genetic research by resolving population structure and speciation events in evolutionary biology, pinpointing sources and transmission chains in epidemiology, and deciphering genotype to phenotype associations in medical and agricultural genomics. Split *k-*mer analysis (SKA)^1^ is a fast and precise method for detecting genetic variation from genomics data, especially single nucleotide polymorphisms (SNPs), without relying on a reference genome. Referred to as “mutation-aware,” split *k-*mers have shown promise for rapid investigations of bacterial outbreaks^2,3^, especially in tracking strain divergence^4^. SKA intuitively adapts *k-*mers by splitting them into three components, a central nucleotide and two flanking sequences of equal length. To ensure specificity, the two flanking sequences must have an exact match in alignments, while to detect SNPs, the central nucleotide(s) is allowed to simultaneously align with mismatches. Eukaryotic mitochondrial genomes share key structural and sequence similarities with bacterial genomes, including their circular organization, gene content, and AT-rich composition, reflecting their evolutionary origin from an ancestral α-proteobacterium^5,6^. We hypothesized split *k*-mers could provide a new, more reproducible method to haplotype mitogenomes as compared to multilocus sequence-based phylogenies (MLST) protocols, which are limited by ambiguous gene selection and alignment trimming (Fig. 1).

**Figure 1.**
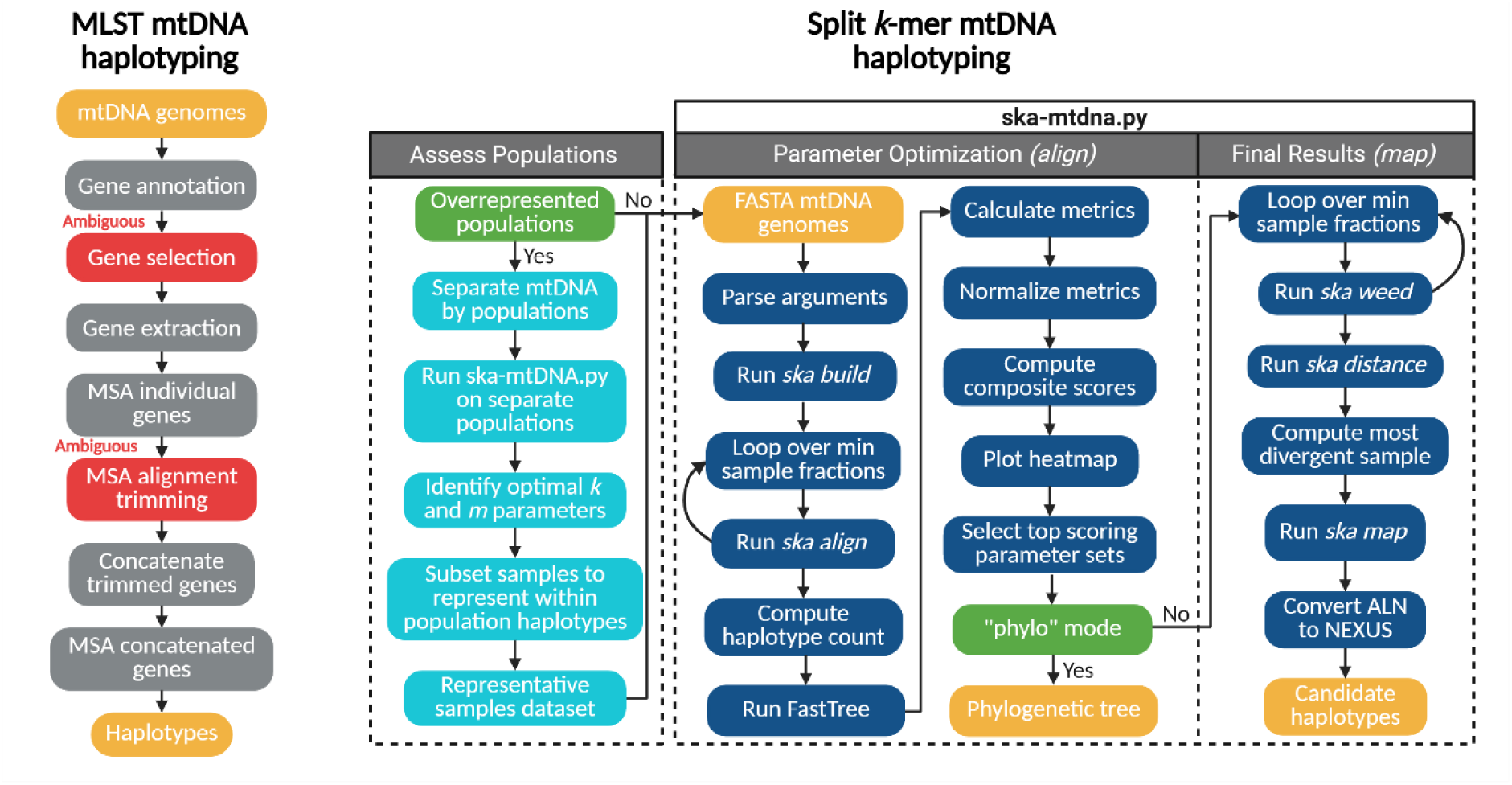
Workflow of MLST mtDNA haplotyping compared to haplotyping mtDNA with split *k-*mers. For MLST genes are annotated, ambiguously selected and trimmed, then concatenated to produce CDS-based haplotypes. The split *k-*mer approach includes two major steps: (1) Populations in a dataset with a fraction of samples greater than the best scoring fraction *m* are haplotyped separately to assign representative samples. (2) Once populations sizes are normalized, ska-mtdna.py is run to explore parameter sets of *k and m* with *ska align* and resulting alignment files are scored. If the default network mode is used, the top scoring *k*-mer lengths of reference-free alignments are refiltered by sample fraction *m* and mapped to the most divergent mitogenome of each parameter set for final haplotyping.

Due to unstandardized gene selection, different gene sets are used for MLST studies of the same species resulting in conflicting evolutionary histories, or incongruence^7^. This produces inconsistent case-specific results that hinder interpretation quality and statistical power when comparing multiple studies, leading to redundancy and a need to reanalyze data^8^. Furthermore, due to varying gene lengths and sequence diversity multiple sequence alignments (MSAs) and whole genome alignments (WGAs) are commonly trimmed to the limiting sequence, removing unreliable regions unique to each study (Fig. 1). Generally, alignment trimming decreases the impact of alignment errors, however this is done at the cost of phylogenetic accuracy^7^. For manually trimmed alignments the exact coordinates of deleted sequence are often not reported making results difficult to reproduce^9^. Programs such as ClipKIT^10^ and CIAlign^9^ aim to resolve ambiguity by removing poorly aligned or highly gapped regions programmatically to enhance phylogenetic accuracy. However, automated MSA trimming is prone to error as excessive trimming may discard valuable phylogenetic information leading to further incongruence^11^. In another case, such as with Gblocks^12^, aggressive trimming can result in entire alignments being removed^10^ impacting a dataset’s representativeness. The inherent issues surrounding MLST studies have shown, for bacterial genomics specifically, there are more accurate methods to construct phylogenies such as SNP calling or the concatenation of homologous sequence^13^, in which the latter is a fundamental concept of SKA.

Segmented *k-*mers have shown promise in assessing whole mitogenome similarity^14^ but have not garnered attention over traditional protocols. Metagenomic sequencing in conjunction with analysis of split *k-*mers may be able to rapidly identify polymorphisms from hundreds of complete mitogenomes at once, an approach that would be scalable to thousands of mitogenomes. Challenges arise when addressing sequence rearrangements in complete mitogenomes; however, we hypothesize that split *k-*mers can effectively handle these rearrangements whether exhibited in protein coding genes, rRNA, or smaller loci encoding tRNA^15–17^.

High-resolution haplotype networks are fundamental to examine populations of individual species while avoiding premature taxonomic grouping from missing data^18^. Haplotyping methods that reduce missing data, detect the greatest number of reliable variants, and result in the accurate reconstruction of population networks provide the greatest haplotype resolution. SKA transparently introduces missing data by assigning ambiguous bases in alignment which are influenced by sample diversity along with two easily assessed parameters; the minimum fraction of samples a *k-*mer must be found in (*m*) and split *k-*mer length (*k*). Sequence conservation of vertically transferred mitochondria allows split *k*-mers to accurately detect haplotypes across multiple generations and populations, although diverse datasets, such as those with different species, can result in excessive ambiguities and loss of data. To ensure the greatest detection of SNPs, without introducing excess ambiguous bases, the effects of both *k* and *m* should be investigated as these parameters are not generalized across different datasets. SKA uses two different methods to produce multiple sequence alignments, *align* and *map.* SKA *align* produces an unordered alignment which is used to build phylograms and haplotype networks; however, to learn where SNPs reside within a mitogenome the *map* algorithm must be used. With ska *map*, split *k-*mers are mapped to a reference for the sole purpose of a coordinate system^2^ and repeated *k*-mers can be masked to avoid inaccuracies when handling short tandem repeats (STRs) in mitochondrial genomes.

In this work, we investigated the ability of SKA to detect mitochondrial haplotypes in three different species: *Diaphorina citri* and *Frankliniella intonsa*, insect vectors of plant pathogens, and *Ovis spp*, the mammalian genus of domesticated sheep. Haplotype results produced with WGA and MLST protocols were compared to haplotype results obtained with SKA2^2^ as follows. Two mitogenome datasets with published MLST haplotype networks from *Diaphorina citri* (n = 31)^19^ and *Fraklienella intonsa* (n = 149)^20^, and one dataset with a WGA haplotype network from *Ovis spp.* (n = 17)^21^ were selected for reanalysis. Lastly, one unpublished dataset of *D. citri* (n = 681) containing citrus grove-collected insect samples from Florida (n = 608), lab colony samples from University of California at Davis in California (n = 5), and lab colony samples from Cornell University in Ithaca, NY (n = 37) combined with available genomes on GenBank was used in this study. *D. citri* is the insect vector of *Candidatus* Liberibacter asiaticus, the invasive, uncultural bacterial pathogen associated with citrus greening disease, currently decimating the Florida citrus industry^22^.

## Results

### SKA mtDNA haplotyping workflow

To streamline the SKA mtDNA workflow we developed a Python script ska-mtdna.py that processes mitogenome FASTA files as input and scores *k* and *m* parameter sets based on alignment statistics. The script is designed to systematically explore combinations of split *k*-mer lengths (*k*) and minimum sample fractions (*m*) to find parameter sets which generate the most accurate and resolute haplotype results (Fig. 1). Lowering the minimum *k*-mer sample fraction filter *m* from default 0.9, which includes more sequence data found in fewer samples, increases haplotype resolution (Fig. 2). In parallel, a decrease in split *k*-mer length *k* increased haplotype resolution by allowing SNPs to be detected closer together. However, the decrease in *k*-mer length and sample fraction filter came at the cost of introducing repeated *k*-mers and ambiguous base pairs which both negatively influence haplotype accuracy when set too low (Fig. 2). Overall, decreasing *k* increased the number of haplotypes, nucleotide diversity, total segregating sites, pairwise nucleotide differences, total nucleotide sites, and haplotype diversity (Fig. 3a). As a relationship between alignment metrics and parameter sets exists, these metrics were implemented in the design of a weighted composite value that robustly scores parameter sets. The optimal *k*-mer lengths we identified were dataset specific, with lengths of 13bp for the *Ovis spp*. dataset, 19bp for the *F. intonsa* dataset, 23bp for *D. citri* GenBank and 17bp for *D. citri* Combined datasets. The varying optimal *k*-mer lengths and sample fractions revealed an importance to assess parameter sets and develop a robust scoring metric to simplify parameter optimization.

**Figure 2.**
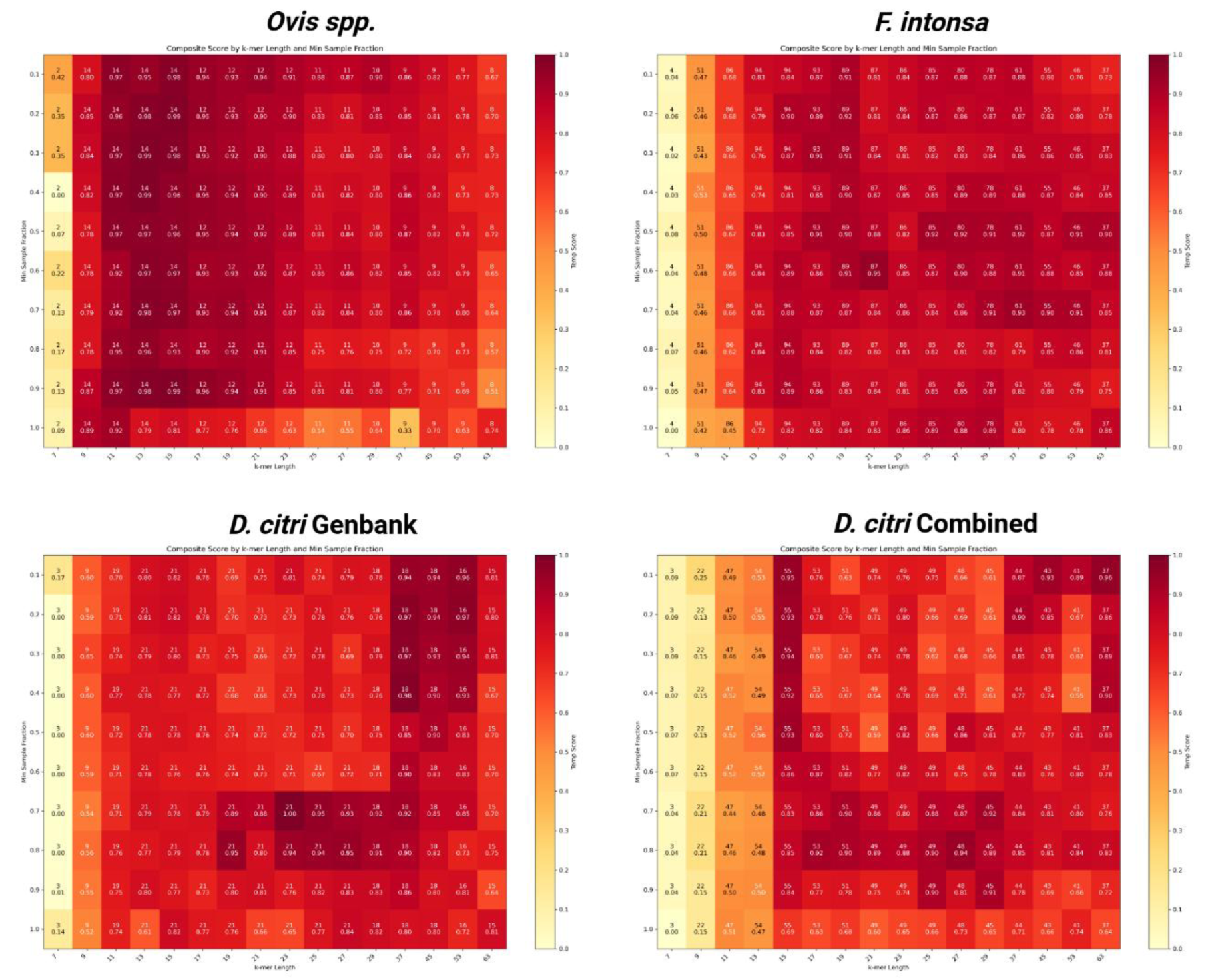
Network mode heatmaps colored by composite score (Temp score) comparing *k*-mer length and minimum sample fractions. Numbers in each cell represent the number of haplotypes detected for each parameter set and the composite score value.

**Figure 3.**
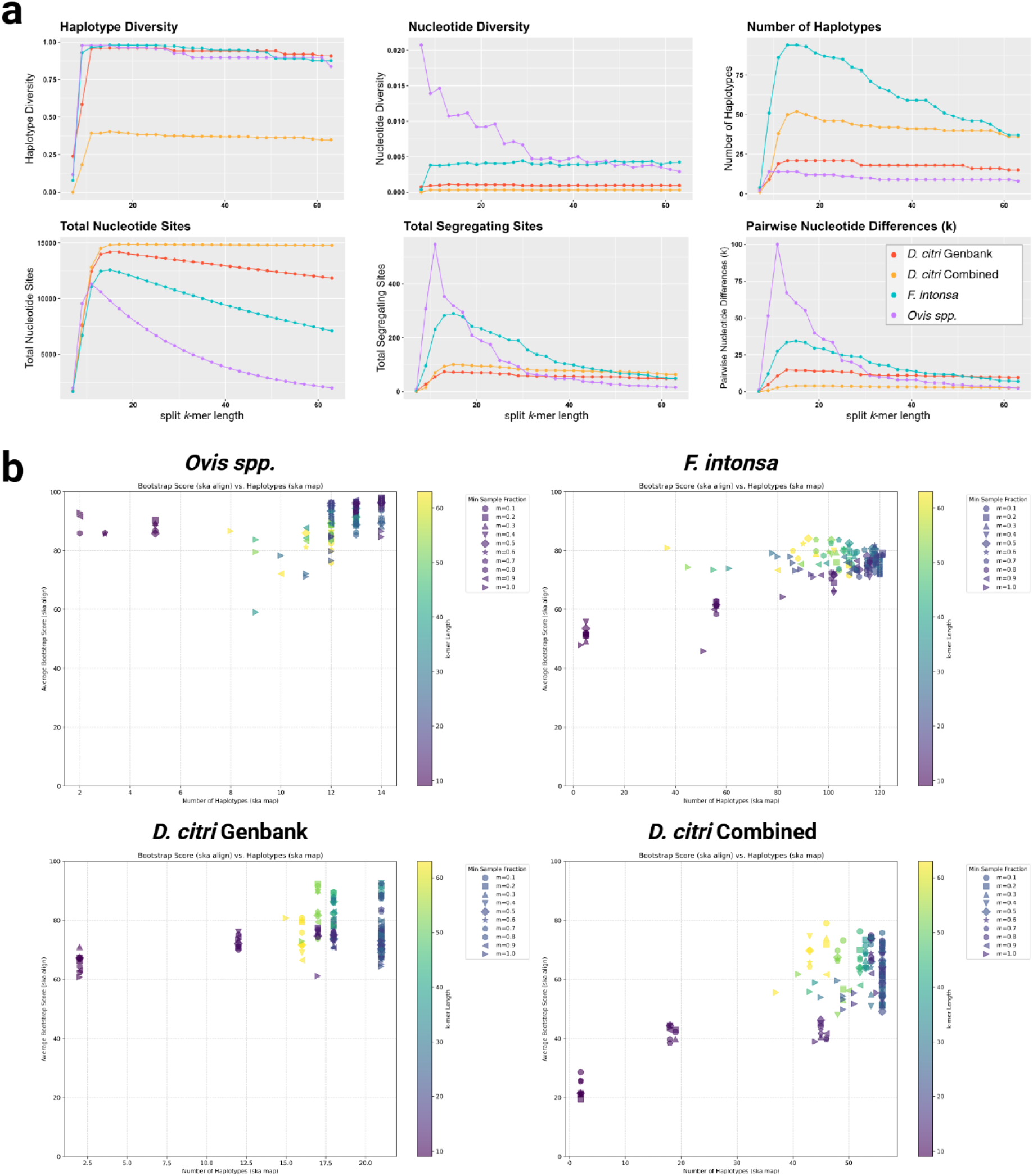
**(a)** Scatterplot depicting the relationships between haplotype count, bootstrap values, *k*-mer length, and sample fraction filters. Red dash lines and red numbers represent the number of haplotypes identified from published analyses using MLST (*F. intonsa* and *D. citri* GenBank) and WGA (*Ovis spp.)*. (**b)** The impact of different *k* values (7 to 63) at *m* = 90 on six statistical measures of population diversity calculated by DnaSP6^25^.

Discrepancies between ska *map* and ska *align* arise from their distinct handling of repeat-associated polymorphisms, with implications for SNP calling accuracy and haplotype inference. We compared the two algorithms for the *D. citri* GenBank dataset and found, for the same parameter sets, the *map* algorithm identifies a greater number of SNPs compared to align (Fig. S1). Upon investigation of the additional SNPs detected we identified “phantom SNPs” that were incorrectly incorporated into alignments for individual samples. Importantly, some of the additional SNPs, but not all, were incorrectly identified due to map mishandling repeated *k*-mers. Polymorphic sites in repeat *k*-mers were marked as ambiguous by the *align* algorithm, however the *map* algorithm integrated these ambiguous polymorphic sites found to be nonexistent in WGA of individual samples. For the *D. citri* GenBank dataset, *k*-mers filtered by a minimum sample fraction of 90% and mapped to the most divergent reference with *map*, resulted in two phantom SNPs two base pairs apart resulting in two new haplotypes (Fig. S1). SKA *align* executed using split *k*-mers filtered at the same minimum sample fraction filter threshold, by either ska *align* or *weed*, marked these variants as ambiguous despite detecting other single-sample SNPs. Inherently ska *align* contains a sample fraction filter as part of the algorithm and ska *weed* is an algorithm independent of sequence alignments used to simply filter *k*-mer profiles. Separate Kalign^23^ WGAs of the two samples with one other sample from the same haplotype identified by *align*, but separated by *map*, revealed no SNPs separating the mitogenomes. Indels only were present in the alignments which did not indicate the absence or addition of the correct nucleotide resulting in the phantom SNPs recorded. When repeats were masked with ska *map*, or the minimum sample fraction was set to 1, the phantom SNPs were set as ambiguous and no longer influenced the final haplotype detection, and yet a greater number of SNPs were still detected compared to *align*. On average, repeat masking removed 5% of SNPs from the most resolute parameters where up to 3% of the SNPs removed were phantom SNPs in STRs. The discrepancy between the two algorithms was further supported by the haplotyping results of the *F. intonsa* dataset (Fig. S2). As a result, the two phases of the ska-mtdna.py script involve a reference-free assessment of *k* and *m* parameter sets ska *align* for initial assessments of *k* and *m* to quickly identify a range of candidate parameter sets, which informs final haplotyping with the *map* algorithm and masked repeats.

The first phase of ska-mtdna.py calculates a composite score for each reference-free alignment by min-max normalizing three metrics - average bootstrap value, number of haplotypes, and valid nucleotide sites - to a value between 0 and 1, then computes a weighted sum using specific weights computed by Sequential Least Squares Programming (SLSQP). The composite score is used to rank *k* and *m* parameters with the goal of finding the most reliable sets for the second phase with the *map* algorithm (Fig 2). The final assessment with map considers the ten highest scoring parameter sets and ranks the *map* results according to haplotype count, number of segregating sites, and lastly by bootstrap scores. The result is a suggested parameter set and nexus files containing ordered alignments. In network mode, haplotype network construction and resolution are prioritized using weights of 0.87 (bootstrap), 0.06 (haplotypes), and 0.07 (valid sites). In phylo mode parameter sets are scored considering the same metrics with an additional weight of -0.1 for gap proportions to support accurate tree construction by avoiding excessively gapped alignments. In both network and phylo modes phylogenetic trees are built with FastTree v2.1^24^ to calculate average bootstrap scores. To identify weights for the robust scoring method of phylograms, we used the metrics of the optimal parameter set identified for a thirty-four mammalian mtDNA dataset^14^ as a target. For each of the five datasets examined, our script accurately identified parameter sets producing the most resolute haplotype network structure, or accurate phylogenetic tree, near identical to published counterparts obtained with MLST or WGA proving its robustness to sample size and diversity.

Reference-based and reference-free approaches to handling masked repeat sequences involve a tradeoff on their use for haplotype network construction or phylogenetic tree analysis. A parallel analysis using a mitogenome dataset from a segmented *k*-mer study^14^ found that split *k*-mers could accurately reconstruct phylogeny of the thirty-four species (Fig. S4) at k = 11 and m = 40. The accuracy of split *k*-mer constructed phylogenetic trees directly correlated to average bootstrap values, which are seen in the associated heatmap, and led to the exploration of relationships between average bootstrap values and metrics indicating accurate haplotype network construction (i.e. total valid nucleotide sites, segregating sites). As bootstrap values are independent of algorithms used to construct haplotype networks, a direct correlation between bootstrap values and optimal haplotyping parameter sets was not identified (Fig. 3b). Yet, average bootstrap values indicate the confidence level of relationships from alignment data and allow for the classification of reliable phylogenies. As multiple parameter sets can produce an identical number of haplotypes (Fig. 3b), bootstrap scoring was used to model the most accurate relationships between the most resolute, near-identical haplotype networks. The differences of gap handling by phylogenetic tree and haplotype network construction algorithms led to the development of two ska-mtdna.py modes, one for haplotype networks and one for phylogenetic trees.

### Comparison with MLST and WGA methods

For the D. citri GenBank dataset SKA improved mitochondrial haplotype resolution (Table 1), providing an unbiased snapshot of where sequence variation occurs across the mitogenome. Split *k-*mers were more sensitive in separating mitochondrial haplotypes of *D. citri* populations with a 24% increase in haplotype resolution compared to a previously published *D. citri* dataset. For the larger and more diverse published datasets of *F. intonsa* and *Ovis spp.*, decreases in haplotype resolution of 2% and 18% were found, respectively (Table 1). We further show that for the *Ovis spp.* dataset, SKA significantly decreased nucleotide diversity (π), and segregating sites (S) compared to WGA, but haplotype count (h), and haplotype diversity (Hd), differences were not significantly different. For the *F. intonsa* dataset SKA significantly decreased π compared to MLST, but h, S, and Hd differences were not significant. Lastly, for the *D. citri* GenBank dataset MLST vs. SKA showed no significant differences in h or Hd, but SKA significantly decreased π compared to MLST, (S untestable due to unreported value). All statistics and *P*-values are reported in Table 1.

**Table 1.** Population statistics of haplotypes represented in Fig. 4 calculated in DNAsp6. **^1^**h = number of haplotypes, ^2^π = nucleotide diversity. ^3^S = number of segregating sites, ^4^Hd = haplotype diversity. Standard error (SE) calculated as the following; h: ^5^SE = √[p * (1 - p) * (1/n₁ + 1/n₂)] where p is a pooled proportion, π: SE = √(Var₁ + Var₂), S: SE = √[p * (1 - p) * (1/L₁ + 1/L₂)], Hd: SE = √(Var₁ + Var₂).

| Dataset | Statistic | WGA | MLST | SKA | n | Length | Difference<br>± SE<sup>5</sup> | P-value |
| --- | --- | --- | --- | --- | --- | --- | --- | --- |
| *Ovis spp.* | h<sup>1</sup> | 17 | - | 14 | 17 | 16463 | 0.1765<br>± 0.0973 | 0.0697 |
| | $\pi^2$ | 0.016 | - | 0.00738 | 17 | 16463 | 0.0086<br>± 0.0001 | 0.0000* |
|  | S<sup>3</sup> | 1387 | - | 451 | 17 | 16463 | 0.0569<br>± 0.0025 | 0.0000* |
|  | Hd<sup>4</sup> | 1 | - | 0.978 | 17 | 16463 | 0.0220<br>± 0.0356 | 0.5363 |
| *F. intonsa* | h | - | 124 | 121 | 149 | 15220 | 0.0201<br>± 0.0443 | 0.6495 |
| | $\pi$ | - | 0.00456 | 0.00352 | 149 | 15220 | 0.0010<br>± 0.0000 | 0.0000* |
|  | S | - | 474 | 433 | 149 | 15220 | 0.0027<br>± 0.0019 | 0.1669 |
|  | Hd | - | 0.997 | 0.994 | 149 | 15220 | 0.0030<br>± 0.0078 | 0.6988 |
| *D. citri*<br>GenBank | h | - | 17 | 21 | 31 | 15038 | -0.1290<br>± 0.1237 | 0.2970 |
| | $\pi$ | - | 0.00099 | 0.00094 | 31 | 15038 | 0.0001<br>± 0.0000 | 0.0026* |
|  | S | - | - | 70 | 31 | 15038 | - | - |
|  | Hd | - | 0.938 | 0.961 | 31 | 15038 | -0.0230<br>± 0.0555 | 0.6788 |

Even though split *k-*mers were less sensitive compared to traditional MLST for *F. intonsa*^20^ and WGA for *Ovis spp.*^21^, split *k*-mers reconstructed each of the five major lineages (L1-L5) of *F. intonsa* as well as species-specific populations of *Ovis spp.* in haplotype networks (Figs. 4a, 4b). Compared to another published haplotype analysis^19^, SKA produced a more resolute haplotype network of globally distributed *D. citri* (Fig. 4c). For the unpublished dataset consisting of 607 mitogenomes, named “*D. citri* Combined”, SKA detected one major haplotype (n = 516) along with other more recently emerging haplotypes within North America (Fig. 4d). Of the North American haplotypes, a lab colony of *D. citri* from Ithaca, NY originating from samples collected in Florida, possessed a unique mtDNA haplotype. Of the samples from California (n = 6), one wild-caught sample, OM181945.1, represented its own haplotype while the other five UC-Davis lab colony samples grouped with the major North American haplotype (Fig. 4d). Mapping Asian *D. citri* haplotype locations based on recorded latitude and longitude^26^ of samples from the *D. citri* GenBank dataset revealed distinct subpopulations of *D. citri* across Southeast Asia (Fig. S5) which further supported the accuracy of split *k*-mer mtDNA haplotyping.

**Figure 4.**
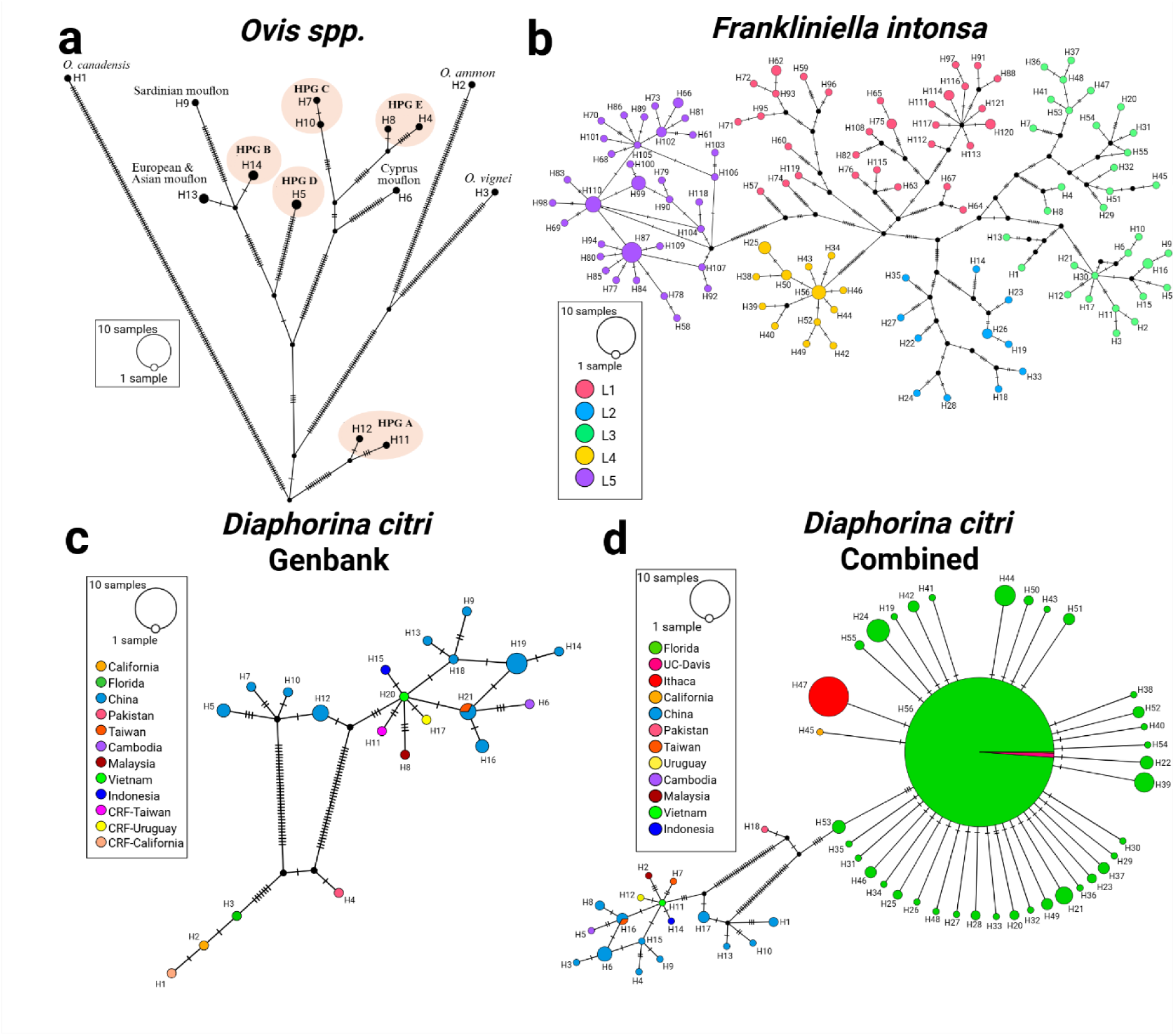
Four haplotype networks obtained using SKA2 *map* and generated in PopART^27^ for three different species. Samples within each haplotype can be found in Tables S1-S4. Haplotypes are ranked from most to least divergent. (**a)** TCS network of 17 samples using *k* = 13 and *m* = 0.1 from the Ovis Genus labeled by known haplogroups^21^. (**b)** Median-joining network using *k* = 19 and *m* = 0.2 of 149 *F. intonsa* mtDNA colored by known taxonomic lineages^20^. (**c)** TCS network using *k* = 23 and *m* = 0.7 of 31 *D. citri* mtDNA. (**d)** TCS network of 681 *D. citri* mtDNA using k = 17 and m = 0.8.

### Reducing missing data and distribution of SNPs across genetic elements

Missing data were exacerbated by increased sample diversity and overrepresented populations which both impacted *k*-mer filtering and SNP detection. The disproportionate representation of a population resulted in the removal of *k*-mers with true SNPs, as those k-mers were not found in a population with a fraction of samples greater than or equal to the minimum sample fraction assigned to *m* (Fig. S3). Therefore, sequences from the same species and equal population representation are important for split *k*-mer mtDNA haplotyping. Datasets of individuals from the same genus may also be investigated using split *k*-mers, although for many genera inherent mtDNA sequence diversity will introduce missing data. Split *k-*mers mapped to the most divergent mitogenome detected SNPs in rRNA, tRNA, CDS, and intergenic sequences proportional to the total amount of sequence each element harbors in a mitogenome, displaying no bias to recombinant prone elements (Fig. 5).

**Figure 5.**
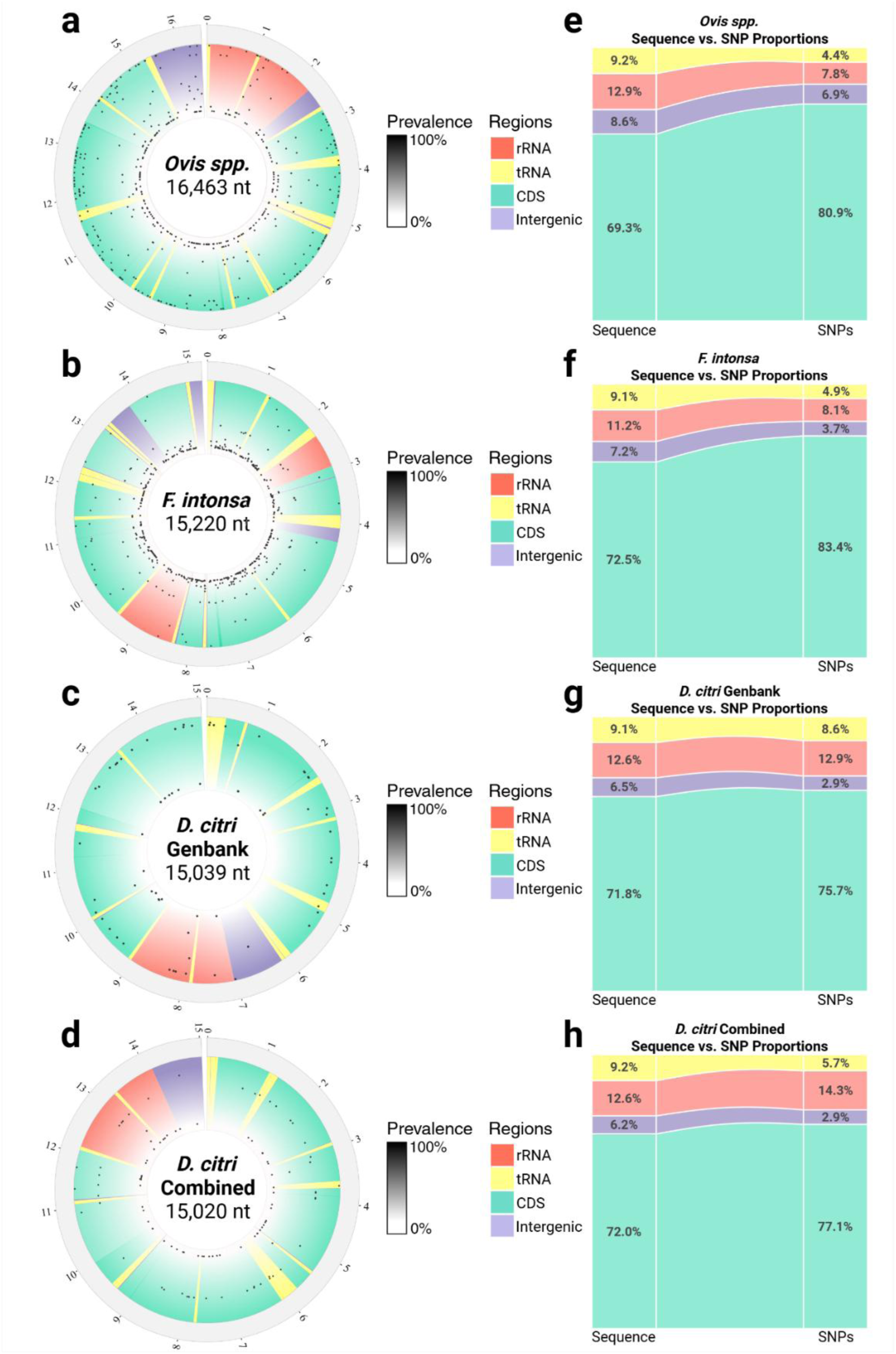
**(a-d)** Distribution of SNPs detected in mitochondrial genomes from four datasets generated with BioCircos^28^. SNPs are ranked by prevalence percentage. Most divergent mitogenomes were used as references. Reference GenBank accessions are as follows; *Ovis spp.*: JN181255.1, *F. intonsa*: OP546405.1, *D. citri* GenBank: OM181945.1, and *D. citri* Combined: MF614822.1. (**e-h)** The percentage of sequence each genetic element occupies in a mitogenome proportional to percentage of SNPs residing in those genetic elements from each dataset. Figures generated in RStudio^29^.

### Epidemiology and genetic associations

We tested whether SKA could detect an association between *D. citri* haplotype and the titer of the citrus greening bacteria, “*Candidatus* Liberibacter asiaticus” (*C*Las) titer within each insect. *C*Las is a circulative, propagative plant pathogen, meaning the *C*Las bacteria replicate within the body of the insect vector during the transmission cycle^4^. Our analysis revealed there are multiple *D. citri* mtDNA haplotypes in Florida and that there is a significant correlation between mtDNA haplotypes of *D. citri* and *C*Las titer for samples collected from the same locations by a Kruskal-Wallis rank-sum test (Fig. 6). Two haplotypes represent individuals with low to no *C*Las titer and four haplotypes represented *D. citri* individuals with high *C*Las titer. Importantly, MLST analysis of 13 concatenated mtDNA protein-coding sequences did not detect the two *D. citri* haplotypes, 39 and 51, which are both associated with *C*Las titer. These data provide evidence for the benefit of split *k*-mers to strengthen epidemiological research in vector borne disease systems.

**Figure 6.**
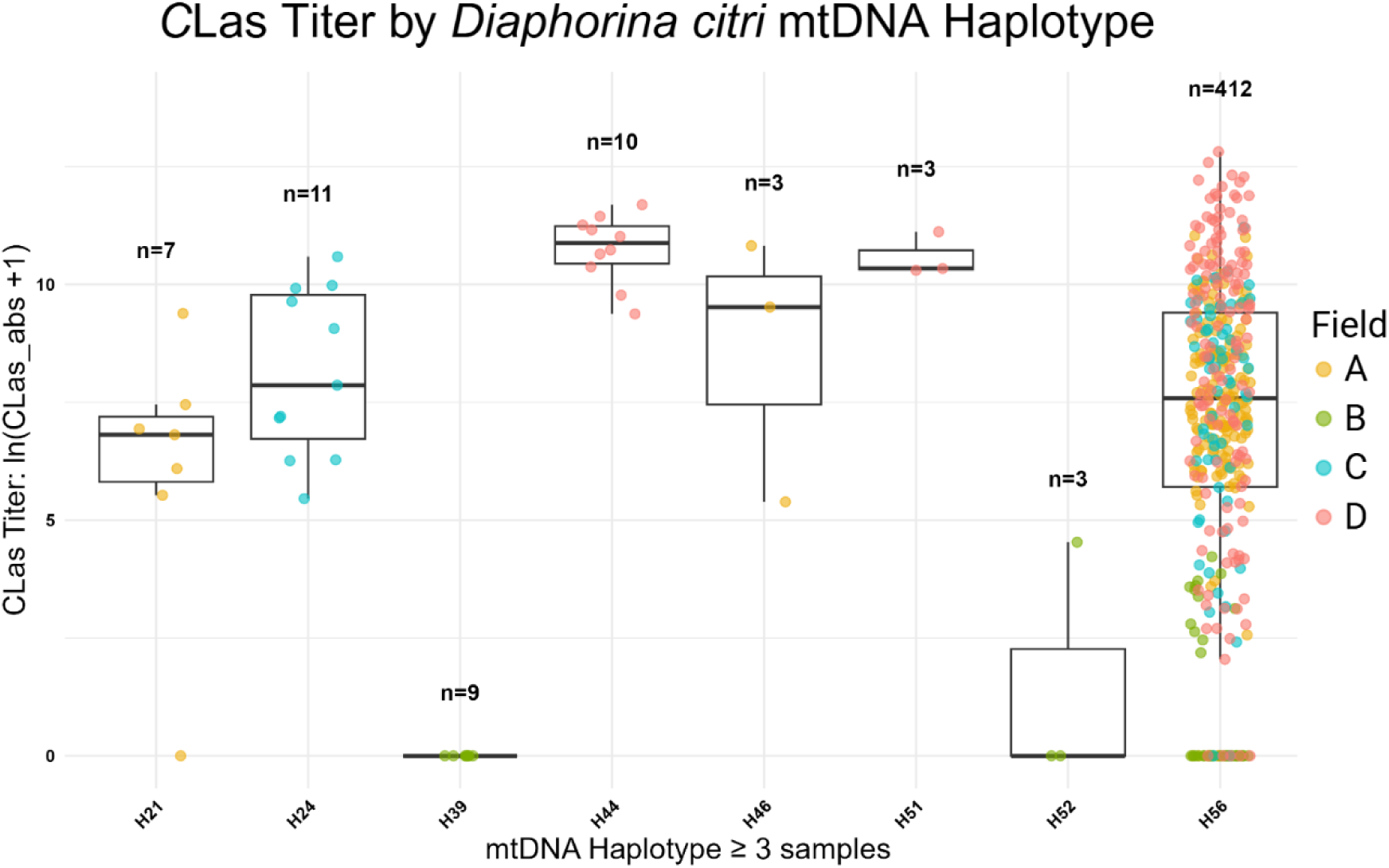
Box plot of titer values per haplotype, with three or more samples, of *D. citri* from Florida. Individual adults were collected from four field sites in Southeast Florida. Field A = Port St. Lucie, Field B = Pioneer, Field C = Whidden Corner, Field D = Montura. A Kruskal-Wallis rank-sum test was performed with *C*Las titer as the dependent variable and haplotype as the independent factor revealed a chi-squared = 52.93 and p-value = 3.826e^-09^ for the 8 haplotypes indicating that *C*Las titer distributions differed significantly across haplotypes. SNPs were located in the following genetic regions for the seven haplotypes identified to have significantly different CLas titers: *H21* – cox2; 3226 TTG>TTT (L>F), *H24* – cox3; 5003 GCA>GTA (A>V), *H39* – 16S rRNA; 12337 T>C, *H44* – atp6; ATA>GTA (I>V), *H46* – cox3; 5341 GGA>AGA (G>R), *H51* – 12S rRNA; 13474 C>T, *H52* – nad1; 11717 ATA->GTA (I>V).

## Discussion

Our analysis aimed to characterize the accuracy of SKA when used as a tool to haplotype mitochondrial genomes. Comparisons of SKA haplotyping results to results obtained with current standard workflows show split *k*-mers generate near-identical or improved haplotype results due to the detection of polymorphisms outside of protein coding sequence. Intergenic sequence has gained interest in genetic research as an increasing number of noncoding DNA functions have been classified. Although, due to the complexity of intergenic sequence and lack of conservation, genetic studies focus mainly on coding sequences where functions are more characterizable. Furthermore, major assumptions underlying phylogenetics are violated if mutations originating from horizonal gene transfer, which are more likely to occur in intergenic sequences, are incorporated into analyses^30^. MLST studies are biased towards polymorphisms residing within conserved protein coding sequences to help avoid this issue despite neglecting polymorphisms in functional intergenic elements and tRNAs. In theory, the use of intergenic sequence for phylogenetics increases the potential introduction of mutations arising from recombination in animal mitogenomes. Albeit, some animal mitochondria, such as higher mammals, rarely experience recombination^31^ and in humans the generational impact of recombinant mtDNA is arguably irrelevant^32^, making split *k-*mers applicable to a more diverse sets of mammalian mitogenomes. Conversely, insects experience high rates of mitochondrial gene rearrangement as a method to correct for high rates of nucleotide substitution^33^. In another study^34^, a greater rate of mitochondrial gene rearrangement in insects was not correlated with a greater nucleotide substitution rate, consistent with the hypothesis of the repair mechanism. Indels, possibly produced from these rearrangements, are not considered in haplotype network construction, allowing split *k-*mers to accurately haplotype whole mitogenomes while handling sequence inversions, duplications, and transpositions.

To simplify the complexity of split *k*-mer mtDNA haplotyping analysis, ska-mtdna.py calculates alignment metrics and bootstrap values with pandas and FastTree, respectively, to score parameter sets. The script essentially automates the complex process of finding optimal SKA2 parameters for mtDNA analysis, balancing multiple quality metrics to identify parameter combinations that produce the most informative alignments for downstream phylogenetic or population genetic analyses. The *align* and *map* algorithms leveraged for haplotype network construction produce similar but different results which highlight ska *map*’s sensitivity to variants found in individual samples and mishandling of ambiguous polymorphisms in repetitive sequence. We hypothesize the distinction between the two algorithms arises from the reference-based *k*-mer coordination where mapped kmers are anchored to the coordinates of a reference sequence allowing for the detection of SNPs in rarer *k*-mers. Identification of a reference sequence is initially carried out with ska *distance* from which an average of all sample pairwise distances are calculated. The sample with the largest average distance is identified as the most divergent and is set as the reference. The *align* reference-free, consensus-based approach failed to identify some individual sample SNPs, likely due to disrupted split *k*-mer continuity and inability to chain non-homologous *k*-mer sequences between samples. The lack of sensitivity to rare SNPs indicated the reference-free approach is limited in detecting haplotypes; however, the *align* algorithm was found to construct more accurate phylogenies, with higher bootstrap values, than the *map* algorithm with masked repeats.

By default, ska-mtdna.py masks repeat *k*-mers to avoid the introduction of phantom SNPs in the final map results, although repeat masking may be turned off. Turning off repeat-mask may include more SNPs within intergenic sequence containing repeat regions at the cost of introducing “phantom SNPs” in alignments, which should be investigated on a sample-to-sample basis. If repeats are not masked, we advise the sample fraction filter is set to 1 as for the *D. citri* datasets an *m* of 0.9 still introduced phantom SNPs into alignments. Excess gaps within STRs, due to masking repetitive *k*-mers, were not present in the reference-free *align* results. As indels are handled differently in haplotype networks versus phylogenetic trees, we determined *align* was most accurate for phylogenetic analysis and *map* was most accurate for haplotype network construction. Haplotype networks, such as those produced by median-joining or TCS algorithms, do not rely on bootstrap resampling and instead are constructed by minimizing mutational steps between haplotypes, often ignoring phylogenetic bootstrap support. Although the algorithms are tailored to separate analyses, we found that average bootstraps are still valuable in selecting the best parameter set from sets identifying the same number of haplotypes and aided in the ultimate selection of high-quality MSAs. For these reasons, our script’s default network mode first identifies candidate parameters for a dataset using the reference-free approach to inform the final, and more biologically informative, analysis with *map*.

Studies examining psyllid mitochondrial haplotypes and vector competence of Liberibacter species are critical to aid in disease epidemiological modeling and to make on-farm management recommendations^35^. Our mtDNA haplotyping of 607 adult *D. citri* mitogenomes revealed a statistically significant correlation between mitochondrial haplotypes and *C*Las abundance in *D. citri* as measured by qPCR. A link between vector haplotype and vectored plant pathogen titer has been reported in other psyllid species. Populations of the potato psyllid, *Batericera cockerelli*, also display distinct haplotypes that are either infected or uninfected with *Candidatus* Liberibacter solanacearum^36,37^. Further research using genome-wide association methods and/or controlled transmission assays with each haplotype-pathogen genotype pair is required to confirm whether these relationships have a genetic^38^ or environmental basis.

Examining large datasets of hundreds of mitogenomes with WGA is memory and time-consuming. Without having to perform read-mapping, gene annotation, multiple sequence alignments, alignment trimming, and gene concatenation split *k*-mer mtDNA haplotyping with ska-mtdna.py provides a straightforward method to haplotype same species mitochondria identical to or better than current standard workflows. The ability of split *k-*mers to haplotype mitogenomes greatly simplifies current protocols and empowers researchers to perform rapid mitogenome haplotyping on small machines with at least 8Gb of memory. While using individuals of the same species, SKA can be a powerful tool, but when examining datasets of more diverse individuals, such as with different species, split *k-*mers should be used with caution and will most likely not produce the most resolute haplotype networks. Yet, we hypothesize that SKA will be a useful tool when examining mitochondrial haplotypes of closely related individuals and will aid researchers in revealing population structure and cryptic species.

Two major limitations to the split *k*-mer approach include the impact of population over representation and sample diversity which both result in the loss of data. The size and diversity of a dataset must be considered as equal population representation and sample diversity are highly influential on the accuracy of the SKA *align* and *map* algorithms. We hypothesize the decrease in haplotype resolution of the *Ovis spp.* dataset is due to the inherent diversity of multiple species causing more SNPs to be spaced closer than the optimal *k*. Regions with multiple SNPs within a subsequence length smaller than k cannot align to other *k*-mers from other samples, leading to their removal. Thus, highly related individuals are best for split *k*-mer mtDNA haplotyping. In the case of *F. intonsa,* the change in haplotype resolution is least significant among the datasets examined (Table 1) where three less haplotypes were detected compared to the published MLST counterpart. As two of the samples with undetected haplotypes belong two *F. intonsa* L5, which contained the most samples, we found the loss of resolution was due to neighboring SNPs. Thrips of the order Thysanoptera consist of multiple cryptic species^39,40^ contributing to complex variation in pathogen vectoring capabilities^41^, a phenomena possibly exacerbated through the use of insecticides and development of insecticide resistance^42,43^. Many lineages of Thrips species are understood to arise from a cryptic species complex resulting in their genetic classifications^44^. As split *k*-mers successfully identified lineages of *F. intonsa* and are most sensitive to mutations in related individuals, the use of split *k*-mers provides a major advantage in the classification of cryptic species, especially for those with cryptic behavior traceable by mitochondrial genome mutations.

This characterization establishes a reproducible and less ambiguous method to haplotype mitochondrial genomes. Future research focused on using SKA in mitochondrial haplotyping will capitalize on the strengths and reduce limitations of the approach. Generally, if more than one third of the total nucleotide sites were removed from mtDNA MSAs after aligning or mapping split *k*-mers with the optimal *k* and *m* parameters, the haplotype structure of the dataset should be examined with other methods such as MLST or WGA. In our experience, generating split *k*-mer MSAs as close to the original length of the mitogenomes being assessed by keeping as many unique *k*-mers as possible, before introducing ambiguous base pairs, is optimal as the latter will hinder haplotype network construction. In our experiments, extensive characterization of the compounding variables (dataset size, population representation, sample diversity, algorithmic discrepancies, and repetitive sequence) was required to create a well-informed and robust workflow. Further optimization of this method is likely required to improve results for more diverse datasets, robust handling of unequal population representation, and to better handle repeated *k*-mers to confidently identify SNPs in repeat sequences

## Conclusion

The use of split *k-*mer analysis sets a foundation for unambiguous and reproducible phylogenetic research of mitochondrial genomes. The ska-mtdna.py script provides robust examination of *k* and *m* parameters using weighted statistics indicative of accurate haplotyping results while preventing excessive missing data. Two major steps – correcting overrepresented populations and optimizing parameters for specific datasets – provide a reproducible method avoiding ambiguity in MLST and WGA alignment trimming. The statistics and parameter sets identified by ska-mtdna.py capture a customizable panel of results to be examined by the end-user and provide guidance to identify quality haplotyping results. By using multiple datasets with varying levels of diversity we show SKA mtDNA haplotyping is robust, provides an unambiguous analysis in parallel to current methods, and can provide higher resolution compared to MLST methods.

## Methods

### *D. citri* insect collection, DNA sequencing, and CLas quantification

Wild-caught *D. citri* samples were collected from Southeast Florida in October of 2018. Extracted DNA was sequenced by Azenta Life Sciences (GENEWIZ) using Illumina paired-end 7x coverage. Methods further describing *D. citri* sample collection and processing are detailed in Mann et al.^45^ *C*Las titer in each insect used in the haplotype association was measured as described in Mann et al.^45^ and distribution of titer among fields was previously reported in the same study.

### Preparation and selection of mitogenome databases

All unpublished *D. citri* mitogenomes were assembled using default metaSPAdes^46^ after quality filtering reads with FaQCs^47^ parameters --mode BWA_plus -q 30 --min_L 50 -n 1 --lc 0.85 --*k-*mer_rarefaction --adapter. Contigs were aligned to the mitogenome reference *Diaphorina citri* isolate FLpsy mitochondrion KY426015.1 with Nucleotide-Nucleotide BLAST 2.16.0+^48^ and the top scoring matches were selected for analysis. *D. citri* mitogenomes comprising less than 97% of the average mitogenome assembly length were considered incomplete and removed from this analysis. All other mitogenomes were downloaded from GenBank^49^ to perform comparative analysis and benchmarking against published research. Datasets selected for reanalysis were required to consist of full mitogenome assemblies and have a published haplotype network. Selections were further narrowed to examine the impact of genus and species level variation, mutation accumulation in mammalian and insect mitogenomes, cryptic species, and dataset size.

### Optimizing *k, m*, and population distribution

To initially capture the effects of split *k-*mer length (*k*) we examined all lengths of odd *k-*mers from 7 to 63 using the default *m* of 90 and ska *align*^2^. Without having to designate a reference, all FASTA alignment files were generated, and the impact of *k* was assessed with population diversity statistics such as the number of haplotypes, haplotype diversity, nucleotide diversity, segregating sites, and pairwise nucleotide differences. Candidate *k-*mer lengths producing the greatest haplotype diversity were then used to assess a lower minimum sample fraction filter (*m*), leading to the inclusion of sequence data found in fewer samples, to review its impact on population diversity statistics. Guided by published results using standardized haplotyping methods, we found optimal parameters sets to be unique to each dataset. Consistent with the theory of reducing missing data, values indicating a high number of segregating sites and total nucleotide sites associated with the most resolute haplotype networks, while large gap proportions indicated excessive missing data and poor haplotype quality.

To provide a more robust analysis and automated workflow we developed the optimize-ska-mtdna.py and ska-mtdna.py scripts with coding assistance from xAI’s Grok Large Language Model. Identification of optimal metric weights was carried out using the Sequential Least Squares Programming (SLSQP) algorithm from SciPy to maximize the composite score of manually identified target parameters. Target parameters were selected by reviewing *map* results, comparing results to published counterparts, and selecting the parameter set which detected the greatest number of haplotypes with strongest bootstrap support.

By default, the ska-mtdna.py script processes FASTA files using SKA2 to generate alignments for various k (7, 9, 11, 13, 15, 17, 19, 21, 23, and 25) and m (0.1 to 1.0 at .1 intervals) values to generate high-quality alignments. Each alignment is validated and metrics including bootstrap scores (via FastTree with 100 replicates), haplotype counts, total valid nucleotide sites (coordinates with only A, C, G, T), segregating sites, and gap proportions are then calculated from the alignment data. Each metric is normalized to a value between 0 and 1 by min-max normalization and is multiplied by weights (wb, wh, wn, ws, wg) to calculate a final composite score. A composite score is calculated as a weighted sum of normalized metrics and scores are saved in initial results alongside a heatmap representing composite score values. SLSQP optimization was used to maximize scoring of target parameter sets producing high quality alignments and networks. The weights used to initialize SLSQP (wb=0.85, wh=0.05, wn=0.1, ws=0.0, wg=0.0) were identified by iteratively testing weight combinations until target parameters were ranked within the top ten scores for each dataset, accounting for randomly seeded bootstrapping based on pseudo-replicates. Initial weights from the reference-free approach were adjusted within preset bounds, prioritizing high bootstrap values (wb=0.8–0.9), while considering lower weights for haplotype count (wh=0.0–0.1), total nucleotide sites (wn =0.05-0.15), segregating sites (0.0-0.05), and gap proportion (-0.05-0). Weights for phylo mode were computed using the same metrics as network mode, but with a different initial weight (-0.05) and bound (-0.15-0) to penalize excessive alignment gap proportions. The objective function maximized the score for each dataset’s target parameter, incorporating penalties for weights that outscore the target parameter and bonuses for weights that increase the composite score of the target parameter.

The number of samples for each population in a multi-population study must be normalized so *k-*mers were not prematurely filtered out by the minimum sample fraction filter. To reduce the size of overrepresented populations, we examined populations individually with SKA *map* after identifying the most divergent haplotype in a population. From the population-specific analysis, one sample from each multi-sample haplotype was selected to represent that haplotype in the complete dataset thus reducing the number of samples from a population, while preserving haplotype resolution. The results of ska-mtdna.py should be used to review individual populations.

To determine if split *k*-mer haplotyping results were significantly different from published MLST or WGA results we determined statistical significance of population metrics. P-values were calculated for the number of haplotypes (h), nucleotide diversity (π), number of segregating sites (S), and haplotype diversity (hd) to determine if split *k*-mer haplotyping significantly increased or decreased these metrics. A P-value ≤ 0.05 was considered statistically significant. For h, we used a two-sample proportion z-test (h/n). For π, we used a Z-test with variance ≈ π / (n(n-1)/2 * L). For S, we used a two-sample proportion z-test (S/L), and for Hd we used a Z-test with variance ≈ Hd(1 - Hd) / n, where n is the number of samples and L is the length of the split *k*-mer alignment.

### Examining the impact of sequence diversity

As sequence diversity is a pitfall of SKA, this study was inherently designed to review the impact of sequence diversity on haplotyping mitochondrial genomes. By including datasets from three species, with differing levels of diversity e.g. different species of the same genus or different lineages of the same species, we not only benchmarked SKA against published datasets but also reviewed how mitogenome diversity decreases SNP detection capabilities of SKA. Thorough exhaustive analysis of each dataset using all *k* lengths and varying *m* proportions captured how these parameters can be adjusted to account for sequence diversity allowing researchers to produce resolute haplotype networks.

### Distribution of SNPs

To assess the distribution of SNPs detected by SKA, we mapped split *k-*mers with optimal values of *k* and *m* for each dataset to the sequence with the longest average predicted genetic distance. The output file in Variant Call Format (VCF) from the ska *map* function was then converted by the snippy^50^ vcf_to_tab tool to produce a tab-delimited input for Biocircos^28^ plotting in R. Custom python scripts were used to calculate both the percentage of SNPs within genetic elements, and the percentage of nucleotides each genetic element harbors within a mitogenomes genome for proportional comparisons.

## Data Availability

All assembled mitogenomes used for this analysis are available on FigShare^51^.

## Code Availability

Custom scripts and code for this analysis are available on GitHub (https://github.com/dougiestuehler/mtDNA_haplotying).

## Acknowledgements

We thank Dr. Steven Higgins and Dr. Marina Mann for their assistance in collecting and processing *Diaphorina citri* samples for Illumina short-read sequencing.

This research is supported by U.S. Department of Agriculture (USDA) Agricultural Research Service Project # 8062-22410-007-000-D and the Emergency Citrus Disease Research and Extension Program (ECDRE) projects 2020-70029-33176 and 2022-70029-038503 from the USDA National Institute of Food and Agriculture. This research was also supported in part by an appointment to the Agricultural Research Service Research Participation Program administered by the Oak Ridge Institute for Science and Education (ORISE) through an interagency agreement between the U.S. Department of Energy (DOE) and the USDA. ORISE is managed by ORAU under DOE contract number DE-SC0014664. All opinions expressed in this paper are the author’s and do not necessarily reflect the policies and views of DOE, or ORAU/ORISE.

## Contributions

Conceptualization: D.S.S. and M.H. Data curation: D.S.S, M.H., L.M.C. Formal analysis: D.S.S. Funding acquisition: M.H. Investigation: D.S.S., M.H. Methodology: D.S.S., M.H. Project administration: M.H., L.M.C. Resources: M.H., L.M.C. Software: D.S.S. Supervision: M.H., L.M.C. Validation: D.S.S., M.H., L.M.C. Visualization: D.S.S., M.H., L.M.C. Writing - original draft: D.S.S and M.H. Writing - review and editing: D.S.S, M.H., L.M.C.

