## Supplementary Data for "Rapid and reproducible haplotyping of complete mitochondrial genomes using split *k-*mers"

**Figure S1.** TCS Haplotype networks of *D. citri* Genbank dataset produced with SKA *align* and SKA *map* algorithms with $k = 15$ and $m = 0.9$, which was the highest scoring parameter set without repeat masking. The *map* algorithm identified two more haplotypes in the dataset (blue and black boxes) showing that the ska *map* function is more sensitive to discern mtDNA haplotypes. The SNP detected with the *map* function separating MF614803.1 and MF614804.1 is 6744A>T in the NADH dehydrogenase subunit 5 gene. The SNP detected with the *map* function separating MF614820.1 and MF614817.1 is 6746T>C in the NADH dehydrogenase subunit 5 gene.

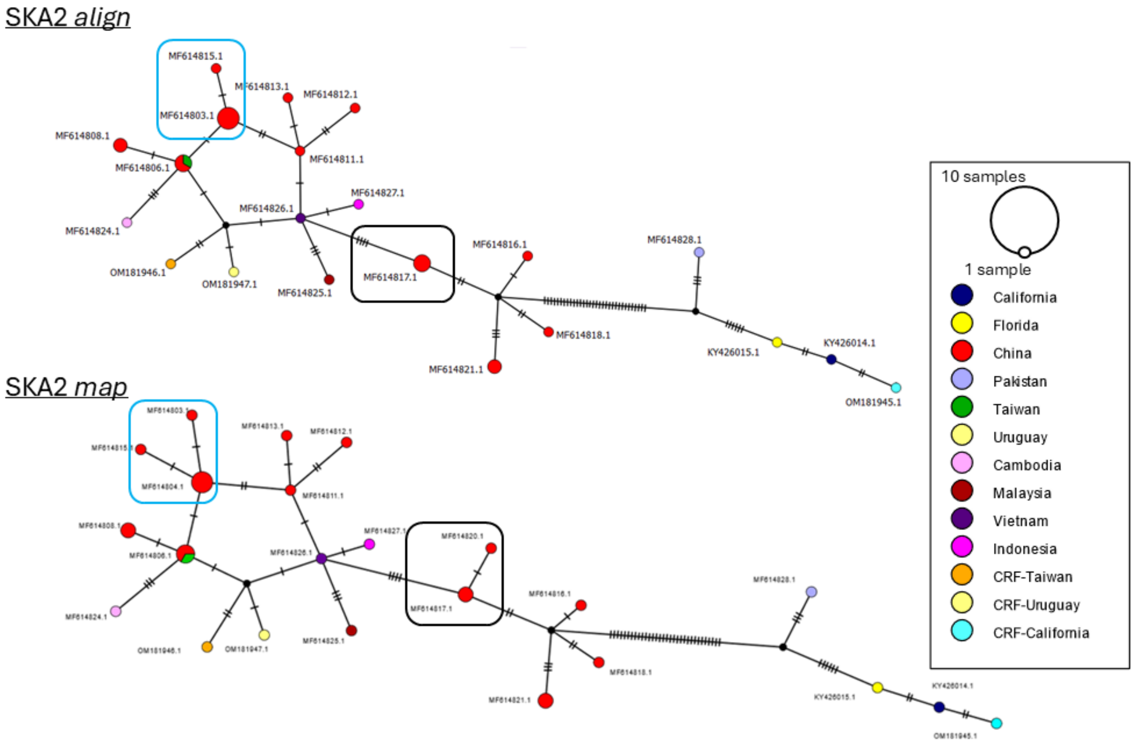

**Figure S2.** Median joining haplotype networks of the *F. intonsa* dataset produced with SKA *align* and SKA *map* algorithms with $k = 19$ and $m = 0.2$. The *map* algorithm identified thirty-two more haplotypes in the dataset showing that the ska *map* function is more sensitive in discerning mtDNA haplotypes.

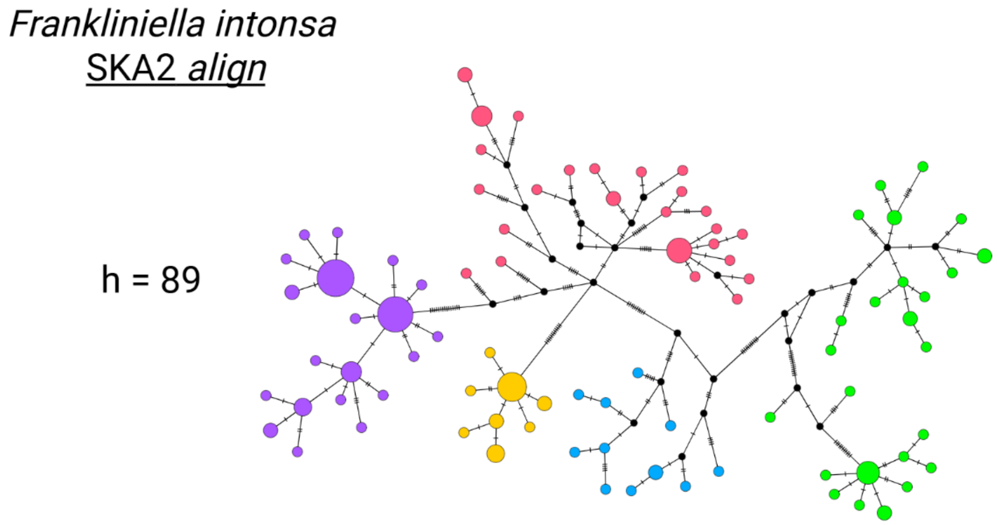


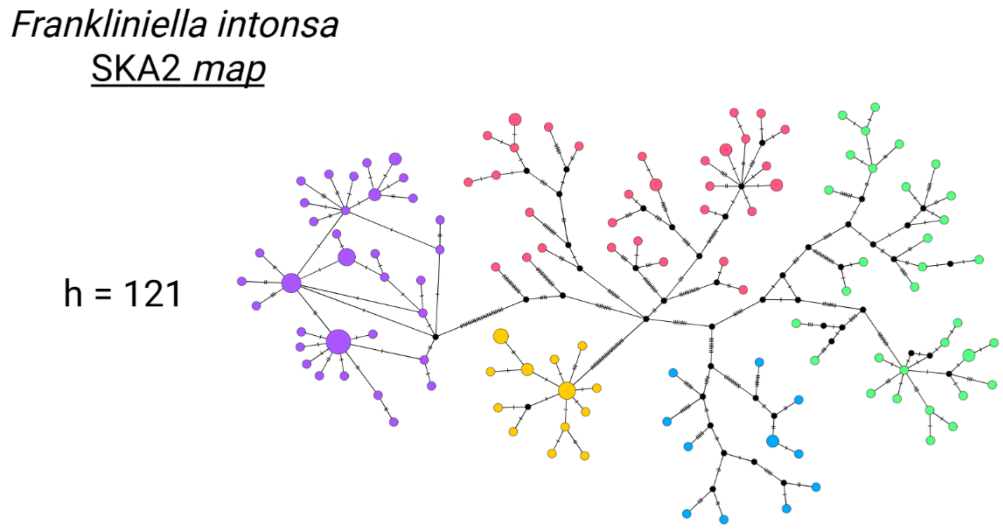

**Figure S3.** Two TCS haplotype networks of the *Diaphorina citri* Combined dataset using $k$ = 15 and $m$ = 90. The top network (a) was made using population assess representative haplotypes and the bottom network (b) used all samples to haplotype. Due to the overabundance of samples from the Florida population, split $k$-mers with SNPs from Asian samples and a California sample were filtered out by the minimum sample fraction filter $m$. This resulted in a lack of haplotype detection and resolution, where four haplotypes of Asian samples were not detected without population assessment and a one-sample haplotype from California was not detected without initial population assessments. After haplotyping the overabundant Florida samples separately using ska *map*, one sample representing each haplotype from the Florida population were refiltered and haplotyped with the rest of the Combined dataset to reduce the impact of population overrepresentation. The $k$-mers unique to a population, containing real SNPs, will not be filtered out as long as the number of samples from that population with the unique $k$-mer/s represent a percentage of the total samples greater than or equal to the minimum sample fraction assigned to $m$.

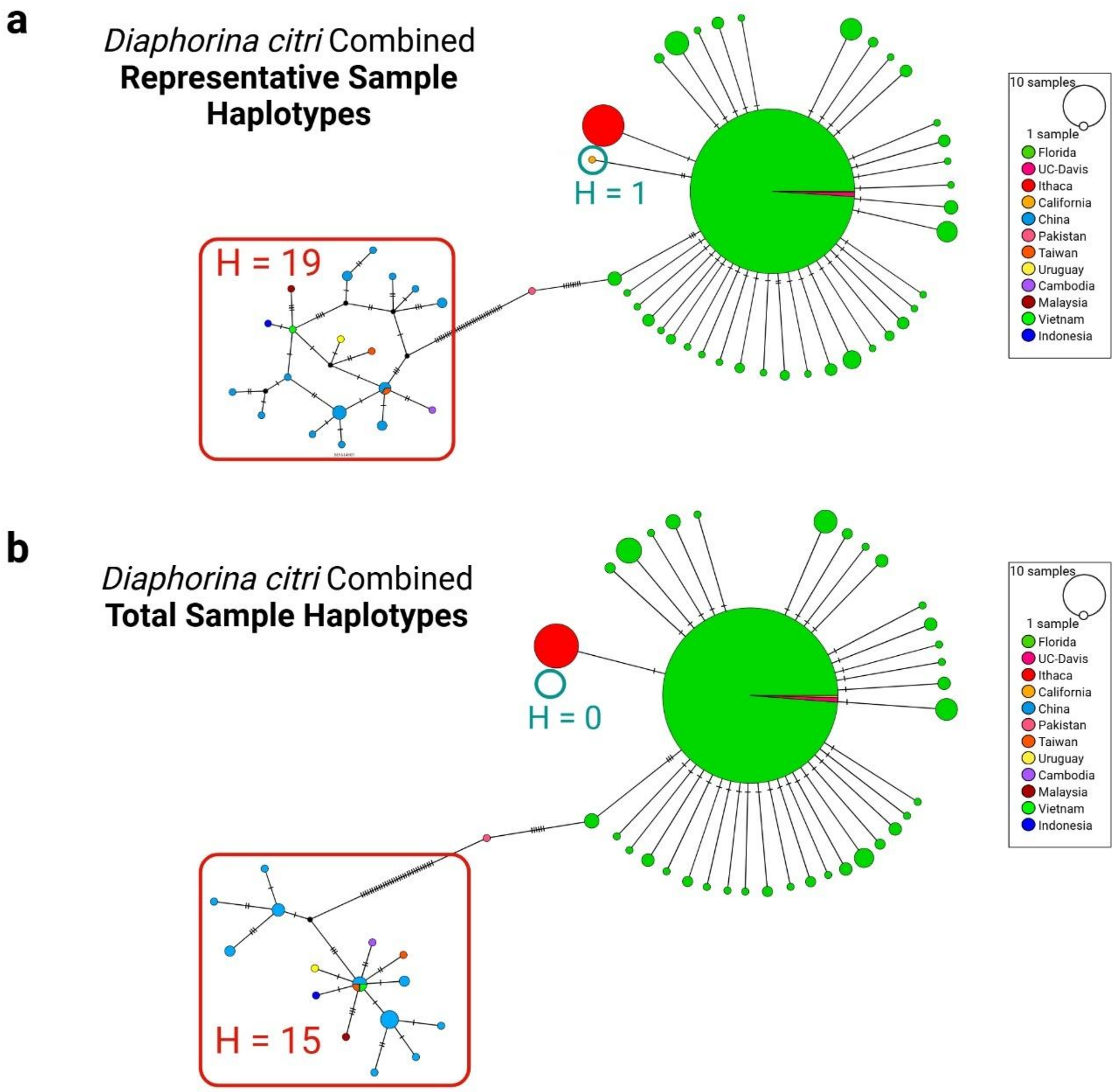

**Figure S4. a.** FastTree phylogenetic tree of 34 mammalian species made with *k* 11 and *m* 40. Tree was built using the optimize-ska-mtDNA.py script where the generalized time-reversible (GTR) model is used. **b.** Heatmap colored by bootstrap support comparing k-mer length and minimum sample fractions. Numbers in each cell are the number of different haplotypes detected in the dataset for each parameter set. **c.** Heatmap of the same structure colored by composite score.

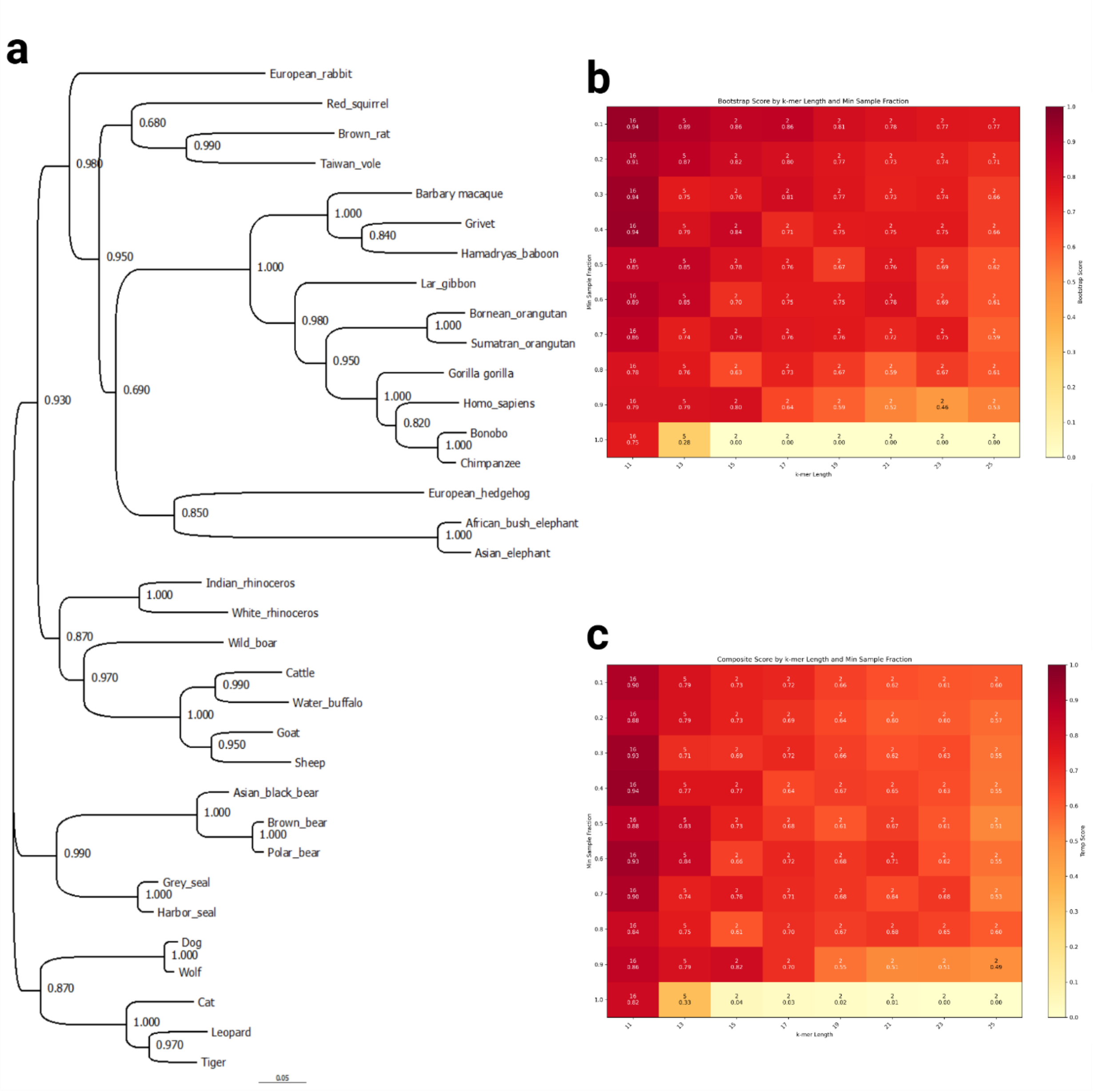

**Figure S5.** TCS haplotype network and map of *D. citri* haplotypes from Southeast Asia produced with the reference free *align* command and dataset from Carlson et al.[16] Haplotypes identified in the TCS network correlate to regional populations of *D. citri* discerning related but distinct haplotypes in the same or different regions.

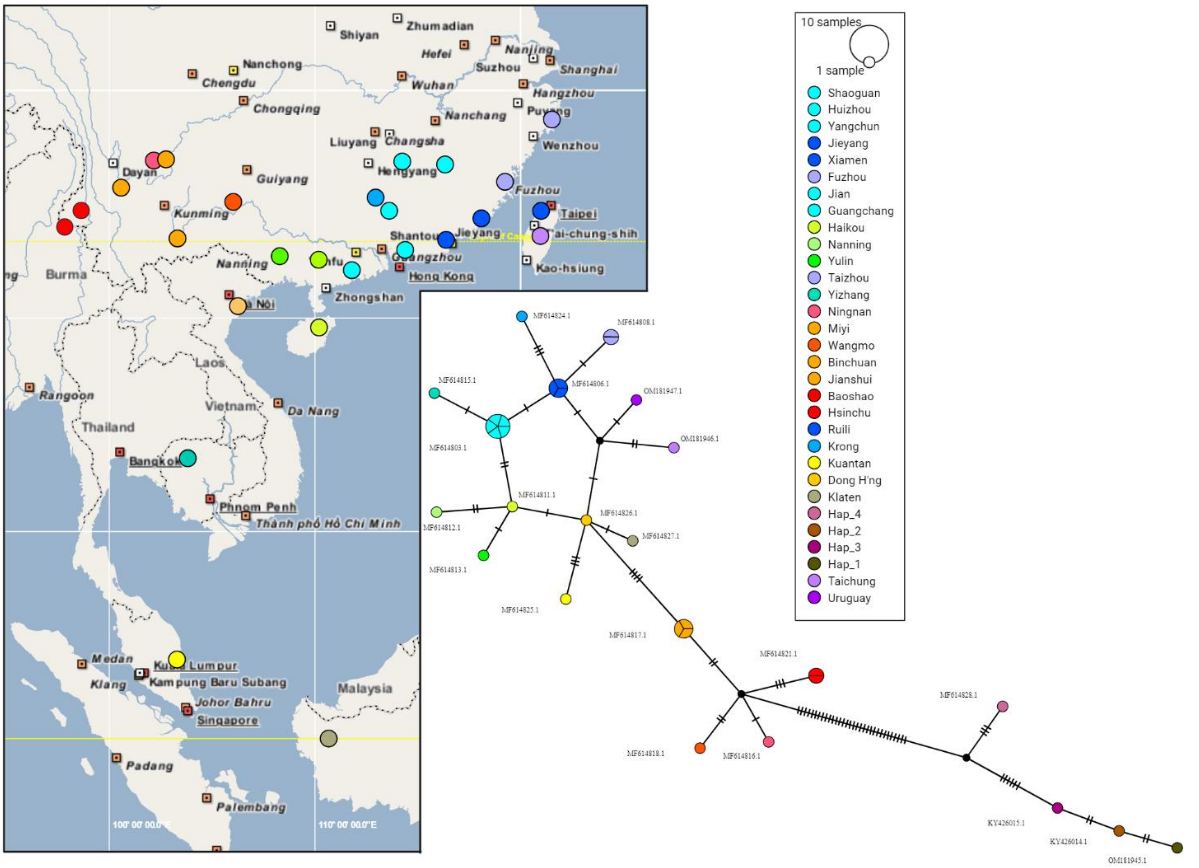

**Table S1.** *Ovis spp.* samples and haplotypes as detected by SKA2 *map.* SNP Distances represent the number of SNPs for that sample found between all samples by pairwise comparisons.

| Haplotype No. | Sample Accessions | SNP Distances |
|---|---|---|
| H1 | JN181255.1 | 393.83 |
| H2 | HM236188.1 | 328.06 |
| H3 | HM236189.1 | 279.16 |
| H4 | HM236183.1 | 172.67 |
| H5 | HM236180.1, HM236181.1 | 171.47 |
| H6 | KF312238.2 | 170.70 |
| H7 | HM236179.1 | 166.77 |
| H8 | HM236182.1 | 166.14 |
| H9 | MG489885.1 | 163.72 |
| H10 | HM236178.1 | 163.55 |
| H11 | HM236174.1 | 158.70 |
| H12 | HM236175.1 | 153.86 |
| H13 | HM236184.1, KF938360.1 | 148.91 |
| H14 | HM236176.1, HM236177.1 | 148.56 |

**Table S2.** *Frankliniella intonsa* samples and haplotypes as detected by SKA2 *map.* SNP Distances represent the number of SNPs for that sample found between all samples by pairwise comparisons.

| Haplotype No. | Sample Accessions | SNP Distances |
|---|---|---|
| H1 | OP546405.1 | 69.57 |
| H2 | OP546494.1 | 68.39 |
| H3 | OP546393.1 | 67.92 |
| H4 | OP546495.1 | 67.83 |
| H5 | OP546399.1 | 67.54 |
| H6 | OP546436.1 | 67.16 |
| H7 | OP546445.1 | 67.16 |
| H8 | OP546492.1 | 66.89 |
| H9 | OP546372.1 | 66.72 |
| H10 | OP546437.1 | 66.69 |
| H11 | OP546368.1 | 66.65 |
| H12 | OP546371.1 | 66.27 |
| H13 | OP546383.1 | 66.22 |
| H14 | OP546395.1 | 66.12 |
| H15 | OP546488.1 | 65.87 |
| H16 | OP546392.1,OP546402.1 | 65.74 |
| H17 | OP546380.1 | 65.69 |
| H18 | OP546459.1 | 65.18 |
| H19 | OP546511.1 | 64.84 |
| H20 | OP546448.1 | 64.83 |

| Haplotype No. | Sample Accessions | SNP Distances |
|---|---|---|
| H21 | OP546431.1 | 64.78 |
| H22 | OP546421.1 | 64.78 |
| H23 | OP546394.1 | 64.67 |
| H24 | OP546463.1 | 64.63 |
| H25 | OP546512.1,OP546513.1,OP546514.1 | 64.11 |
| H26 | OP546508.1,OP546396.1 | 64.09 |
| H27 | OP546477.1 | 64.05 |
| H28 | OP546374.1 | 64.02 |
| H29 | OP546479.1 | 63.95 |
| H30 | OP546433.1 | 63.77 |
| H31 | OP546468.1 | 63.28 |
| H32 | OP546446.1 | 63.05 |
| H33 | OP546452.1 | 62.95 |
| H34 | OP546455.1 | 62.85 |
| H35 | OP546470.1 | 62.83 |
| H36 | OP546466.1 | 62.83 |
| H37 | OP546375.1 | 62.74 |
| H38 | OP546498.1 | 62.66 |
| H39 | OP546515.1 | 62.6 |
| H40 | OP546516.1 | 62.49 |
| H41 | OP546420.1 | 62.38 |
| H42 | OP546500.1 | 62.3 |
| H43 | OP546475.1 | 62.2 |
| H44 | OP546471.1 | 62.18 |
| H45 | OP546442.1 | 62.04 |
| H46 | OP546460.1 | 61.72 |
| H47 | OP546462.1 | 61.37 |
| H48 | OP546478.1 | 61.25 |
| H49 | OP546486.1 | 60.92 |
| H50 | OP546429.1,OP546427.1 | 60.91 |
| H51 | OP546472.1 | 60.9 |
| H52 | OP546482.1 | 60.83 |
| H53 | OP546476.1 | 60.75 |
| H54 | OP546458.1 | 60.66 |
| H55 | OP546501.1 | 60.59 |
| H56 | OP546474.1,OP546376.1,OP546453.1,OP546481.1 | 60.37 |
| H57 | OP546398.1 | 58.35 |
| H58 | OP546435.1 | 58.14 |
| H59 | OP546469.1 | 57.67 |
| H60 | OP546381.1 | 57.38 |
| H61 | OP546401.1 | 57.35 |
| H62 | OP546424.1,OP546423.1 | 57.05 |
| H63 | OP546507.1 | 56.96 |
| H64 | OP546373.1 | 56.91 |
| H65 | OP546480.1 | 56.9 |

| Haplotype No. | Sample Accessions | SNP Distances |
| --- | --- | --- |
| H66 | OP546430.1,OP546426.1 | 56.69 |
| H67 | OP546497.1 | 56.56 |
| H68 | OP546408.1 | 56.5 |
| H69 | OP546415.1 | 56.48 |
| H70 | OP546391.1 | 56.44 |
| H71 | OP546499.1 | 56.19 |
| H72 | OP546422.1 | 56.14 |
| H73 | OP546441.1 | 56.12 |
| H74 | OP546510.1 | 56.11 |
| H75 | OP546443.1,OP546485.1 | 56.02 |
| H76 | OP546509.1 | 56.02 |
| H77 | OP546447.1 | 55.88 |
| H78 | OP546406.1 | 55.85 |
| H79 | OP546491.1 | 55.84 |
| H80 | OP546418.1 | 55.78 |
| H81 | OP546384.1 | 55.69 |
| H82 | OP546465.1 | 55.67 |
| H83 | OP546490.1 | 55.49 |
| H84 | OP546411.1 | 55.4 |
| H85 | OP546417.1 | 55.34 |
| H86 | OP546449.1 | 55.33 |
| H87 | OP546504.1,OP546410.1,OP546432.1,OP546434.1,OP546439.1,OP546440.1,OP546451.1,OP546502.1 | 55.29 |
| H88 | OP546378.1 | 55.28 |
| H89 | OP546400.1 | 55.12 |
| H90 | OP546505.1 | 55.05 |
| H91 | OP546379.1 | 55.02 |
| H92 | OP546416.1 | 54.97 |
| H93 | OP546444.1 | 54.96 |
| H94 | OP546386.1 | 54.94 |
| H95 | OP546454.1 | 54.9 |
| H96 | OP546457.1 | 54.82 |
| H97 | OP546489.1 | 54.58 |
| H98 | OP546390.1 | 54.56 |
| H99 | OP546503.1,OP546409.1,OP546412.1,OP546487.1 | 54.53 |
| H100 | OP546385.1 | 54.5 |
| H101 | OP546388.1 | 54.41 |
| H102 | OP546407.1,OP546403.1 | 54.31 |
| H103 | OP546370.1 | 54.28 |
| H104 | OP546382.1 | 54.21 |
| H105 | OP546450.1 | 54.2 |
| H106 | OP546404.1 | 53.91 |
| H107 | OP546413.1 | 53.91 |
| H108 | OP546461.1 | 53.83 |
| H109 | OP546419.1 | 53.51 |

| Haplotype No. | Sample Accessions | SNP Distances |
|---|---|---|
| H110 | OP546387.1,OP546369.1,OP546389.1,OP546438.1,OP546506.1 | 53.49 |
| H111 | OP546483.1 | 53.39 |
| H112 | OP546414.1 | 53.35 |
| H113 | OP546467.1 | 53.08 |
| H114 | OP546464.1,OP546428.1 | 53.05 |
| H115 | OP546397.1 | 53.05 |
| H116 | OP546493.1 | 53.02 |
| H117 | OP546425.1 | 52.82 |
| H118 | OP546496.1 | 52.4 |
| H119 | OP546484.1 | 52.39 |
| H120 | OP546473.1,OP546377.1 | 52.02 |
| H121 | OP546456.1 | 51.82 |

**Table S3.** *Diaphorina citri* Genbank samples and haplotypes as detected by SKA2 *map*. SNP Distances represent the number of SNPs for that sample found between all samples by pairwise comparisons.

| Haplotype No. | Sample Accessions | SNP Distances |
|---|---|---|
| H1 | OM181945.1 | 46.43 |
| H2 | KY426014.1 | 45.47 |
| H3 | KY426015.1 | 45.35 |
| H4 | MF614828.1 | 37.48 |
| H5 | MF614821.1, MF614822.1 | 14.38 |
| H6 | MF614824.1 | 13.05 |
| H7 | MF614818.1 | 12.62 |
| H8 | MF614825.1 | 12.6 |
| H9 | MF614812.1 | 12.07 |
| H10 | MF614816.1 | 12.02 |
| H11 | OM181946.1 | 11.5 |
| H12 | MF614817.1, MF614819.1, MF614820.1 | 11.17 |
| H13 | MF614813.1 | 11.1 |
| H14 | MF614815.1 | 10.98 |
| H15 | MF614827.1 | 10.67 |
| H16 | MF614808.1, MF614814.1 | 10.48 |
| H17 | OM181947.1 | 10.23 |
| H18 | MF614811.1 | 10.13 |
| H19 | MF614803.1,MF614804.1, MF614805.1, MF614809.1, MF614810.1 | 10.02 |
| H20 | MF614826.1 | 9.7 |
| H21 | MF614806.1, MF614807.1, MF614823.1 | 9.58 |

**Table S4.** *Diaphorina citri* Combined samples and haplotypes as detected by SKA2 *map.* SNP Distances represent the number of SNPs for that sample found between all samples by pairwise comparisons.

| Haplotype No. | Sample Accessions | SNP Distances |
|---|---|---|
| H1 | MF614822.1,MF614821.1 | 37.37 |
| H2 | MF614825.1 | 37.19 |
| H3 | MF614815.1 | 37.02 |
| H4 | MF614812.1 | 36.95 |
| H5 | MF614824.1 | 36.95 |
| H6 | MF614803.1,MF614804.1,MF614805.1,MF614809.1, MF614810.1 | 36.89 |
| H7 | OM181946.1 | 36.7 |
| H8 | MF614814.1,MF614808.1 | 36.7 |
| H9 | MF614813.1 | 35.98 |
| H10 | MF614818.1 | 35.94 |
| H11 | MF614826.1 | 35.77 |
| H12 | OM181947.1 | 35.72 |
| H13 | MF614816.1 | 35.7 |
| H14 | MF614827.1 | 35.22 |
| H15 | MF614811.1 | 35 |
| H16 | MF614806.1,MF614807.1,MF614823.1 | 34.23 |
| H17 | MF614819.1,MF614820.1,MF614817.1 | 33.69 |
| H18 | MF614828.1 | 24.54 |
| H19 | CA-G3 | 23.64 |
| H20 | 29,ACP29 | 23.35 |
| H21 | AB-A12,AC-C3,AC-D2,AD-A11,AD-A12,AD-B7,AD-C7 | 22.92 |
| H22 | 14,4,5,BA-B4 | 22.44 |
| H23 | 38,ACP38 | 22.44 |
| H24 | 65,CA-B8,CA-C7,CA-D9,CA-E5,CA-F2,CA-F5,CA-F9,CA-H2,CA-H7,CA-H8,CA-H9 | 22.44 |
| H25 | AA-A1,AA-F4 | 22.44 |
| H26 | AA-B6 | 22.44 |
| H27 | AA-D4 | 22.44 |
| H28 | AA-F11,AB-F1 | 22.44 |
| H29 | AA-F9 | 22.44 |
| H30 | AB-A8 | 22.44 |
| H31 | AB-B9 | 22.44 |
| H32 | AC-C12 | 22.44 |
| H33 | AC-E3 | 22.44 |
| H34 | AC-F3 | 22.44 |
| H35 | AC-G5 | 22.44 |
| H36 | AD-B4 | 22.44 |
| H37 | AD-B9,AD-E5,AD-F3 | 22.44 |
| H38 | B27 | 22.44 |

| Haplotype No. | Sample Accessions | SNP Distances |
|---|---|---|
| H39 | BA-A11,BA-B11,BA-C8,BA-D10,BA-D7,BA-E8,BA-F11,BA-F8,BA-G10 | 22.44 |
| H40 | BA-B2 | 22.44 |
| H41 | CA-D4 | 22.44 |
| H42 | D12,D21,D22 | 22.44 |
| H43 | DA-G3 | 22.44 |
| H44 | DB-C1,DB-D4,DB-E1,DB-E7,DB-E8,DB-F2,DB-F4,DB-G7,DB-G9,DB-H7 | 22.44 |
| H45 | OM181945.1 | 22.44 |
| H46 | AC-C5,AC-H4,AD-F11 | 22.43 |
| H47 | 19E,20E,22E,24E,2E,3E,4E,7E,9E,A5,E5A,E5B,E5E,E6,L2,SE14,SE15,SE18,SE19,SE20,SE4,SE6,SE8,T11,T12,T13,T14,T15,T16,T3,T4,T5,T6,T9,V1,V12,V2 | 22.41 |
| H48 | AA-C9 | 22.38 |
| H49 | 28,AC-B3,ACP28 | 22.36 |
| H50 | 96,DA-F4 | 22.36 |
| H51 | DA-C2,DA-D1,DA-H3 | 22.36 |
| H52 | BA-A1,BA-F1,BA-H2 | 22.31 |
| H53 | AB-C8,AB-G5,B14,B21 | 22.08 |
| H54 | BA-G4 | 21.99 |
| H55 | CA-C10,CA-D10 | 21.91 |
| H56 | 13,16,18,2,20,21,22,26,27,3,33,36,39,40,46,47,49,50,51,52,53,54,55,56,57,58,59,6,60,61,64,66,68,70,71,72,73,74,76,77,78,79,80,81,82,83,85,86,87,88,89,9,90,91,93,94,95,A21,A23,A24,AA-A12,AA-A2,AA-A4,AA-B1,AA-B11,AA-B12,AA-B2,AA-B4,AA-B7,AA-B9,AA-C1,AA-C12,AA-C4,AA-C5,AA-D12,AA-D8,AA-E1,AA-E12,AA-E9,AA-F1,AA-F12,AA-G11,AA-G12,AA-G5,AA-G6,AA-G7,AA-G9,AA-H1,AA-H11,AA-H3,AA-H4,AA-H7,AA-H8,AA-H9,AB-A10,AB-A11,AB-A3,AB-A5,AB-A6,AB-A7,AB-B1,AB-B10,AB-B12,AB-B4,AB-B5,AB-B6,AB-B7,AB-B8,AB-C10,AB-C5,AB-C6,AB-C9,AB-D10,AB-D12,AB-D2,AB-D5,AB-D6,AB-D8,AB-E1,AB-E10,AB-E11,AB-E2,AB-E7,AB-E9,AB-F10,AB-F11,AB-F12,AB-F3,AB-F5,AB-F9,AB-G1,AB-G12,AB-G6,AB-G8,AB-G9,AB-H10,AB-H2,AB-H5,AB-H6,AB-H7,AB-H9,AC-A10,AC-A11,AC-A12,AC-A3,AC-A4,AC-A5,AC-A6,AC-A7,AC-B1,AC-B11,AC-B2,AC-B4,AC-B6,AC-B7,AC-B8,AC-B9,AC-C11,AC-C4,AC-C8,AC-C9,AC-D1,AC-D11,AC-D3,AC-D4,AC-D5,AC-D7,AC-D8,AC-D9,AC-E10,AC-E11,AC-E12,AC-E2,AC-E4,AC-E5,AC-E6,AC-E7,AC-E8,AC-F4,AC-F5,AC-F8,AC-G1,AC-G10,AC-G11,AC-G12,AC-G3,AC-G9,AC-H1,AC-H2,AC-H3,AC-H6,AC-H8,AC-H9,ACP25,ACP26,ACP27,ACP33,ACP36,ACP39,A | 21.45 |

| Haplotype No. | Sample Accessions | SNP Distances |
| --- | --- | --- |
| | CP40,ACP46,ACP47,ACPA18,ACPA21,ACPA23,ACPA24,AD-A1,AD-A2,AD-A6,AD-B1,AD-B12,AD-B2,AD-B3,AD-B5,AD-B6,AD-C1,AD-C2,AD-C3,AD-C4,AD-C5,AD-C6,AD-C8,AD-D10,AD-D12,AD-D5,AD-D6,AD-D7,AD-E10,AD-E12,AD-E4,AD-E9,AD-F10,AD-F12,AD-F2,AD-F4,AD-G1,AD-G11,AD-G2,AD-G3,AD-G4,AD-G5,AD-G8,B1,B12,B13,B24,B3,B7,BA-A10,BA-A12,BA-A4,BA-A8,BA-A9,BA-B10,BA-B3,BA-B5,BA-B8,BA-B9,BA-C10,BA-C11,BA-C12,BA-C4,BA-C5,BA-C6,BA-C9,BA-D12,BA-D6,BA-D8,BA-D9,BA-E10,BA-E11,BA-E12,BA-E2,BA-E4,BA-E5,BA-E7,BA-E9,BA-F10,BA-F12,BA-F5,BA-F6,BA-F9,BA-G12,BA-G2,BA-G3,BA-G9,BA-H1,BA-H3,BA-H4,BA-H5,BA-H8,BA-H9,C1,C10,C11,C17,C21,C22,C23,C24,C3,C7,C8,CA-A1,CA-A10,CA-A11,CA-A12,CA-A3,CA-A4,CA-A5,CA-A6,CA-A7,CA-A9,CA-B1,CA-B11,CA-B12,CA-B2,CA-B3,CA-B4,CA-B5,CA-B6,CA-B7,CA-B9,CA-C1,CA-C11,CA-C12,CA-C2,CA-C3,CA-C4,CA-C5,CA-C8,CA-D1,CA-D11,CA-D12,CA-D2,CA-D3,CA-D6,CA-D7,CA-D8,CA-E1,CA-E10,CA-E11,CA-E12,CA-E3,CA-E4,CA-E7,CA-E8,CA-F1,CA-F3,CA-F4,CA-G1,CA-G10,CA-G11,CA-G12,CA-G2,CA-G4,CA-G5,CA-G6,CA-G7,CA-G9,CA-H1,CA-H10,CA-H11,CA-H3,CA-H4,CA-H5,CA-H6,D14,D23,D24,D6,D9,DA-A1,DA-A10,DA-A11,DA-A12,DA-A3,DA-A4,DA-A5,DA-A6,DA-A7,DA-A8,DA-A9,DA-B1,DA-B10,DA-B11,DA-B2,DA-B3,DA-B5,DA-B6,DA-B7,DA-B8,DA-B9,DA-C1,DA-C10,DA-C11,DA-C12,DA-C3,DA-C4,DA-C5,DA-C6,DA-C9,DA-D10,DA-D12,DA-D2,DA-D3,DA-D4,DA-D5,DA-D6,DA-D7,DA-D8,DA-E1,DA-E10,DA-E11,DA-E12,DA-E2,DA-E4,DA-E7,DA-E8,DA-E9,DA-F10,DA-F11,DA-F12,DA-F2,DA-F3,DA-F5,DA-F6,DA-F7,DA-F8,DA-F9,DA-G1,DA-G10,DA-G11,DA-G12,DA-G2,DA-G5,DA-G6,DA-G8,DA-G9,DA-H1,DA-H10,DA-H11,DA-H2,DA-H4,DA-H5,DA-H6,DA-H8,DA-H9,DB-A1,DB-A10,DB-A11,DB-A12,DB-A2,DB-A3,DB-A5,DB-A6,DB-A8,DB-A9,DB-B1,DB-B11,DB-B12,DB-B2,DB-B3,DB-B4,DB-B5,DB-B6,DB-B7,DB-B8,DB-B9,DB-C10,DB-C11,DB-C12,DB-C2,DB-C3,DB-C5,DB-C6,DB-D1,DB-D10,DB-D12,DB-D2,DB-D3,DB-D5,DB-D9,DB-E10,DB-E11,DB-E2,DB-E3,DB-E4,DB-E6,DB-E9,DB-F10,DB-F11,DB-F12,DB-F3,DB-F5,DB-F6,DB-F7,DB-F8,DB-G1,DB-G10,DB-G11,DB-G2,DB-G3,DB-G4,DB-G5,DB- | |

| Haplotype No. | Sample Accessions | SNP Distances |
|---|---|---|
| | G6,DB-G8,DB-H10,DB-H12,DB-H2,DB-H4,DB-H5,DB-H6,DB-H8,DB-H9,H1,H2,H3,I3,I9,KY426014.1,KY426015.1 | |
